# A Computational Approach to Design Potential siRNA Molecules as a Prospective Tool for Silencing Nucleocapsid Phosphoprotein and Surface Glycoprotein Gene of SARS-CoV-2

**DOI:** 10.1101/2020.04.10.036335

**Authors:** Umar Faruq Chowdhury, Mohammad Umer Sharif Shohan, Kazi Injamamul Hoque, Mirza Ashikul Beg, Mohammad Kawsar Sharif Siam, Mohammad Ali Moni

**Affiliations:** Department of Biochemistry and Molecular Biology, University of Dhaka, Bangladesh; Department of Genetic Engineering and Biotechnology, Univeristy of Dhaka, Bangladesh; Department of Pharmacy, Brac University, 66 Mohakhali, Dhaka 1212, Bangladesh; WHO Collaborating Centre for eHealth, School of Public Health and Community Medicine, Faculty of Medicine, University of New South Wales (UNSW), Sydney, Australia

**Keywords:** SARS-CoV-2, Nucleocapsid phosphoprotein, Surface glycoprotein, siRNA, siDirect

## Abstract

An outbreak, caused by a RNA virus, SARS-CoV-2 named COVID-19 has become pandemic with a magnitude which is daunting to all public health institutions in the absence of specific antiviral treatment. Surface glycoprotein and nucleocapsid phosphoprotein are two important proteins of this virus facilitating its entry into host cell and genome replication. Small interfering RNA (siRNA) is a prospective tool of the RNA interference (RNAi) pathway for the control of human viral infections by suppressing viral gene expression through hybridization and neutralization of target complementary mRNA. So, in this study, the power of RNA interference technology was harnessed to develop siRNA molecules against specific target genes namely, nucleocapsid phosphoprotein gene and surface glycoprotein gene. Conserved sequence from 139 SARS-CoV-2 strains from around the globe was collected to construct 78 siRNA that can inactivate nucleocapsid phosphoprotein and surface glycoprotein genes. Finally, based on GC content, free energy of folding, free energy of binding, melting temperature and efficacy prediction process 8 siRNA molecules were selected which are proposed to exerts the best action. These predicted siRNAs should effectively silence the genes of SARS-CoV-2 during siRNA mediated treatment assisting in the response against SARS-CoV-2

## 1. Introduction

COVID-19, A pandemic affecting lives of billions of people worldwide, has confronted humanity in the commencement of 2020, is caused by a viral pathogen, severe acute respiratory syndrome coronavirus 2 (SARS-CoV-2) or 2019-nCoV Initial symptoms of this disease mainly include fever, cough, fatigue, dyspnea & headache[1,2] or it may be asymptomatic[3]. The spike glycoprotein of SARS-CoV-2 binds directly with the surface cell angiotensin converting enzyme II (ACE2) receptor present on alveolar epithelial cells of lung facilitating virus entry, replication and triggers cytokine cascade mechanism[4]. In severe cases, patient may die due to massive alveolar damage and progressive respiratory failure[1,5]. The current detection process of SARS-CoV-2 carried out by most countries is using real-time RT-PCR, although several other methods are also being developed[6–8]. Incubation period for the virus ranges between 2–14 days [9] and in some cases, transmission is also reported during asymptomatic period[10]. Some recent studies suggest that bats are likely reservoir hosts for SARS-Cov-2 but the identity of the intermediate host that might have facilitated transfer to human still remain elusive with some studies indicating pangolins [11]. SARS-CoV-2 is assumed to spread mainly from person-to-person through respiratory droplets produced when an infected person sneezes and coughs or between people who are in close contact[5].

Coronaviruses are genetically classified into four main genera: Alphacoronavirus, Betacoronavirus, Gammacoronavirus, and Deltacoronavirus[12]. The first two genera generally infect mammals, while the last two mostly cause disease in birds. The genome size of coronaviruses ranges between approximately 26-32 kb and includes about 6 to 11 open reading frames (ORFs)[13]. Nucleocapsid protein (N), small envelope protein (E), spike surface glycoprotein (S) and matrix protein (M) are the four major structural proteins of coronavirus and all of which are essential to produce a structurally complete virus[14,15]. The nucleocapsid protein (N) is a multifunctional protein comprising three distinct and highly conserved domains: two structural and independently folded structural regions, namely the N terminal domain and C-terminal domain, which are separated by a intrinsically disordered RNA-binding domain[16]. The primary role of CoV N protein is to package the genomic viral genome into flexible, long, helical ribonucleoprotein (RNP) complexes called nucleocapsids[17]. Apart from these, N protein is essential for viral assembly, envelope formation, genomic RNA synthesis, cell cycle regulation and viral pathogenesis [18–20], Spike glycoprotein (S) is a viral fusion protein which forms homotrimers protruding from the viral surface[21] and mediates virus entry to cell[22]. S contains two functional subunits: S1 & S2 subunits. The S1 subunit includes the receptor-binding domain(s) and contributes to stabilization of the membrane-anchored S2 subunit that contains the fusion machinery [23]. As the coronavirus S glycoprotein is surface-exposed and mediates entry into host cells and N nucleocapsid protein are essential for genome replication, these could be the main targets for designing therapeutics[24].

Silencing of mRNA or post-transcriptional gene silencing by RNA interference (RNAi) is a regulatory cellular mechanism. RNAi is a prospective tool for the control of human viral infections [25–27]. Small interfering RNAs (siRNAs) and micro RNAs (miRNAs) are involved in the RNA interference (RNAi) pathway, where they hybridize to complementary mRNA molecules and neutralizes mRNA causing suppression of gene expression or translation[28]. Studies show that, combinations of chemically synthesized siRNA duplexes targeting genomic RNA of SARS-CoV results in therapeutic activity of up to 80% inhibition [29]. siRNAs directed against Spike sequences and the 3’-UTR can inhibit the replication of SARS-CoV

As of April 10, total infected case is 1,615,059 and among these patients 96,791 people have died which means case fatality rate (CFR) is approximately 5.99%. The alarming phenomenon is the exponential growth of total infection case and death number (https://www.worldometers.info/coronavirus/). Treatment of this increased number of people is not possible as no antiviral drug is still available specifically for SARS-CoV-2 and there is a lack on appropriate medical response. In silico approaches are a general trend to discover novel therapeutic approaches [30–33] and for the viruses there is no exception to this [34].Therefore, in this study, we have designed siRNAs specific to various conserved region of nucleocapsid phosphoprotein & surface glycoprotein genes of SARS-CoV-2 and finally predicted 8 universal siRNA molecules against nucleoprotein and glycoprotein genes which will inhibit the translations of these proteins and allow the host to discard this infection. siRNAs are designed against both nucleoprotein and glycoprotein as both are important for the survival of virus[22,35] and targeting these proteins may cause viral inhibition[29,36].

## 2. Materials and methods

### 2.1. Sequence retrieval from NCBI

Coding sequences from 139 genomes of Severe acute respiratory syndrome coronavirus 2 (SARS-CoV-2) were retrieved from NCBI Virus [37] portal (https://www.ncbi.nlm.nih.gov/labs/virus/vssi/#/) (Supplementary Table 1). Nucleocapsid phosphoprotein and surface glycoprotein sequences were manually extracted and curated from the retrieved data using bash scripting in linux computer platform.

### 2.2. Multiple sequence alignment & phylogenetic tree construction

ClustalW [38] algorithm was employed to perform multiple sequence alignment. Maximum likelihood phylogenetic trees were constructed with a bootstrap value of 500. Tamura Nei [39] model of evolution was selected while constructing the phylogenetic tree. MEGA-X [40] and MEGA-CC [41] programs were was used for alignment formation, phylogenetic tree construction. iTOL online tool [42] (https://itol.embl.de/) was used in order to visualize the phylogenetic trees.

### 2.3. Target recognition & siRNA designing

Target-specific siRNAs were designed with the help of siDirect web server [43]. Rules of Ui-Tei [44], Amarzguioui [45] and Reynolds [46] were used (Table 1) and the melting temperature was kept below 21.5 °C as a default parameter for siRNA duplex.

**Table 1.**
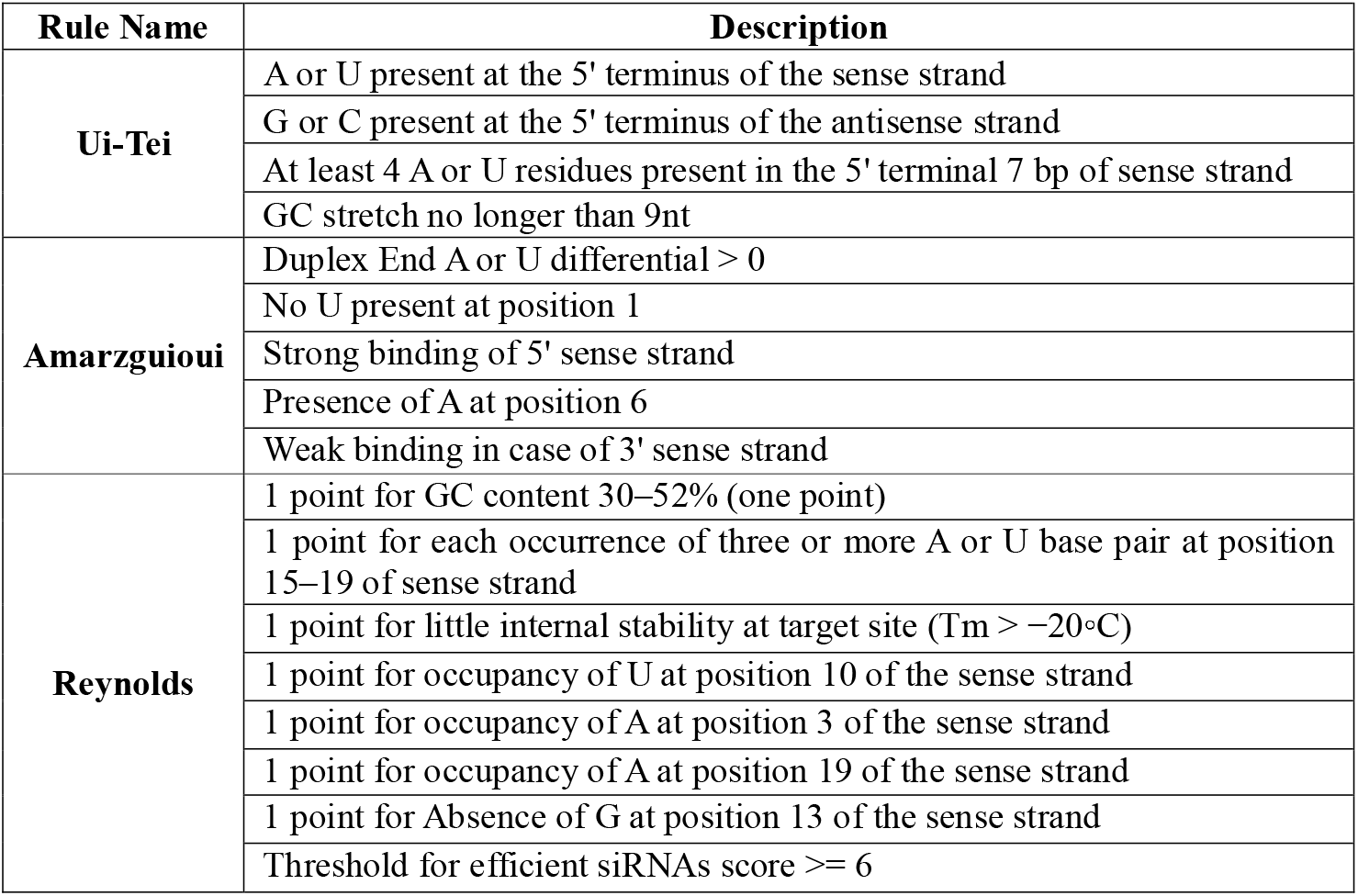
Rules/Algorithm to construct siRNAs.

### 2.4. Off target similarity search using BLAST

Blast search was performed against human genome and transcriptome using the standalone blast package [47] to identify the possible off target matches. The e-value was set to 1e-10 to reduce the stringency of the search condition thereby increasing the chances of random matches.

### 2.5. GC content calculation & secondary structure prediction

OligoCalc [48] was used to calculate the GC content. The secondary structure of siRNAs were predicted along with the respective free energy using MaxExpect [49] program in the RNA structure web server [50]. The higher values indicate better candidates as those molecules are less prone to folding.

### 2.6. Computation of RNA-RNA interaction through thermodynamics

Higher interaction between the target and the guide strand serves a better predictor for siRNA efficacy. Therefore, the thermodynamic interaction between the target strand and the siRNA guide strand was predicted with the aid of DuplexFold [51] program of the RNA structure web server [50].

### 2.7. Computation of heat capacity & concentration plot

DINA Melt webserver [52] was used to generate heat capacity and concentration plot. The ensemble heat capacity (Cp) is plotted as a function of temperature, with the melting temperature Tm (Cp) (Supplementary Table 6 & Supplementary table 7). The contributions of each species to the ensemble heat capacity shown by detailed heat capacity plot. Also, the point at which the concentration of double-stranded molecules of one-half of its maximum value defines the melting temperature Tm(Conc) was shown using the concentration plot-Tm (Conc).

### 2.8. Predicted siRNA Validation

siRNAPred server (http://crdd.osdd.net/raghava/sirnapred/index.html) was used to validate the predicted siRNA species. The predicted siRNAs were evaluated against the Main21 dataset using support vector machine algorithm and the binary pattern prediction approach. siRNAPred score greater than on 1 predicts very high efficacy, score ranging 0.8-0.9 predicts high efficacy and score ranging 0.7-0.8, predicts moderate efficacy. In total, 78 siRNAs were used for efficacy prediction.

## 3. Results

### 3.1. Evolutionary divergence analysis shows conserved pattern between strains

Phylogenetic tree was constructed using 139 sequences for both nucleocapsid phosphoprotein and surface glycoprotein separately. Only a handful number of sequences showed significant divergence (bootstrap value > 60%) (Fig 2, Supplementary Table 8). This suggest that most of the viral sequences have been conserved sequences and therefore used to construct siRNA which will cover a wide range of SARS-CoV-2 strain.

**Fig. 1.**
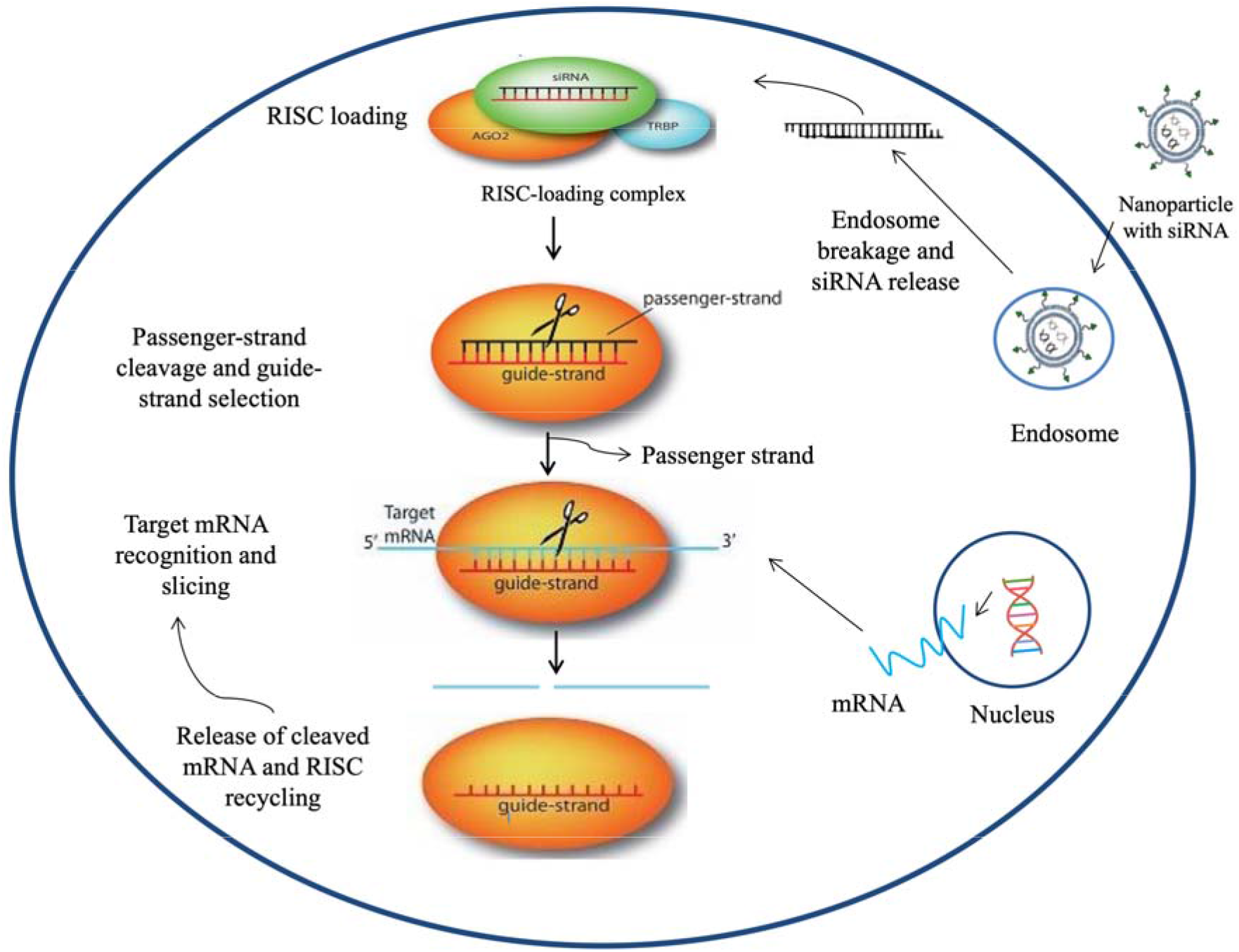
Graphical representation of the siRNA-mediated gene silencing mechanism. Nanoparticles loaded with siRNA are taken up through endocytosis by the cells. These particles are then trapped into the endosomes. siRNA escape endosomes and release siRNA into the cytoplasm due to pH responsive mechanism or proton sponge effect. Once generated, siRNA is loaded into RNA-indiced silencing complex comprising of RNA-binding protein TRBP and Argonaute (AGO2). AGO2 opts the siRNA guide strand, then excises and ejects the passenger strand. After that, the guide strand pairs with its complementary target mRNA and AGO2 slices the target. After slicing, the cleaved target mRNA is released and RISC is recycled for another few rounds of slicing using the same guide strand [55].

**Fig. 2.**
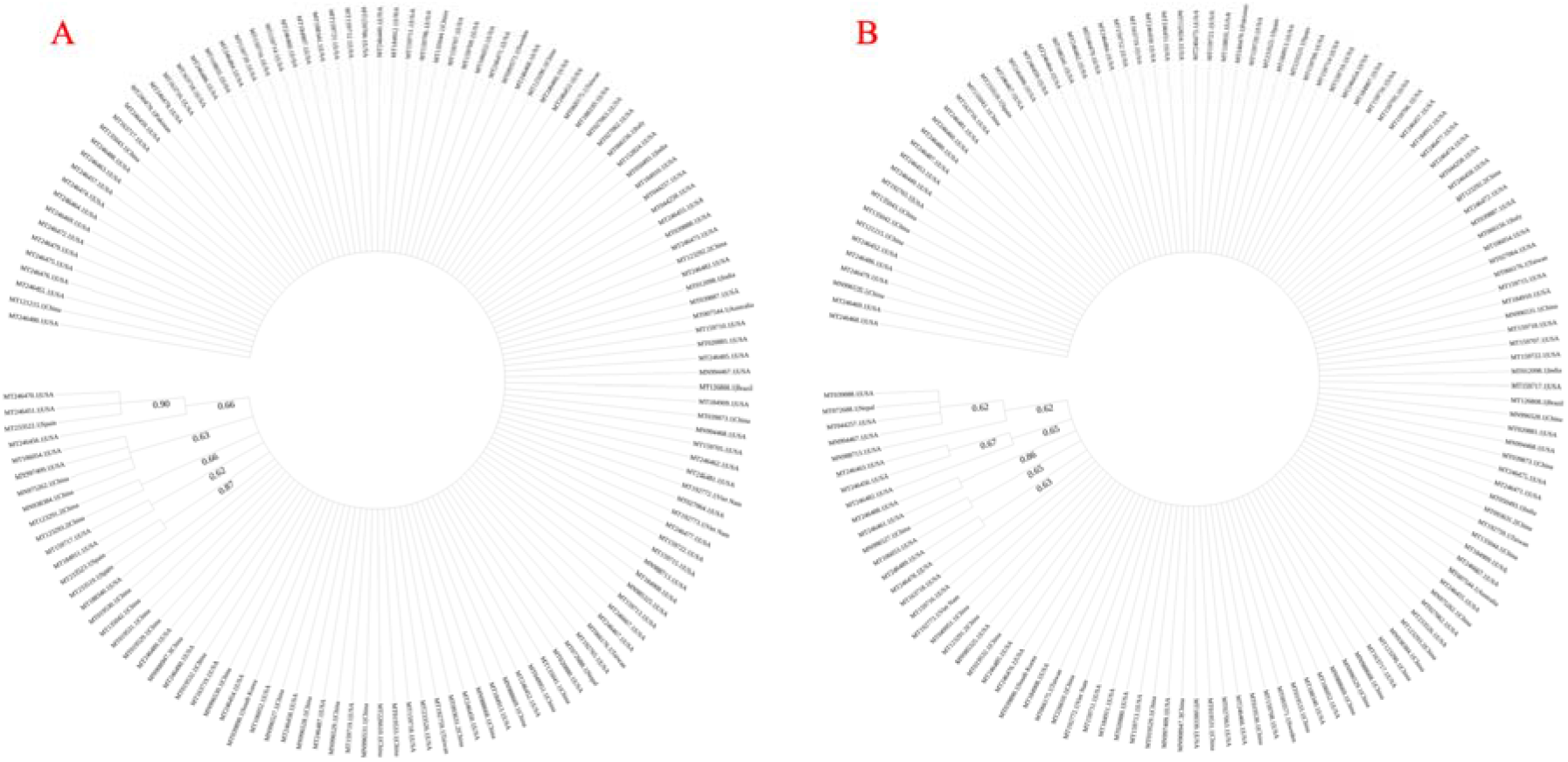
Radial phylogenetic tree of A. nucleocapsid phosphoprotein and B. surface glycoprotein using 139 strains of SARS-CoV-2 from around the world. The bootstrap value for tree construction was set to 500 and Tamura-Nei model of evolution was used for both trees.

### 3.2. siDirect predicted 78 siRNA

siDirect web server predicted 8 siRNA for nucleocapsid phosphoprotein and 70 siRNA for surface glycoprotein (Supplementary Table 4 & Supplementary table 5) that maintains all the parameters. Seed target duplex stability (Tm) values for all the predicted siRNAs were less than 21.5 °C which suggests the ability of predicted siRNAs to avoid non-target binding.

### 3.3 Off-target binding exclusion using blast

Standalone blast [47] search against human genome and transcriptome did not reveal any off-target match. This shows that our predicted siRNA would not interact in any other places other than the viral target location.

### 3.4. GC content calculation & secondary structure determination

GC content analysis of the predicted siRNAs were ranged 31% to 43% for nucleocapsid phosphoprotein (Supplementary Table 6) and 10% to 40% for surface glycoprotein (Supplementary Table 7). Molecules that have GC content below 31.6% were eliminated. Also, the calculated free energy of folding ranged from 1.4 to 2 for nucleocapsid phosphoprotein (Supplementary Table 6) and from 1.3 to 2 for surface glycoprotein (Supplementary Table 7). The associated secondary structures were also determined.

### 3.5. Thermodynamics of target-guide strand interaction

Free energy of binding between target and guide strand were calculated. The values spanned from −35.8 to −31 for nucleocapsid phosphoprotein (Supplementary Table 6) and −36.6 to −21.6 for surface glycoprotein (Supplementary Table 7).

### 3.6. Heat capacity & duplex concentration plot determination

The Tm(Cp) and Tm(Conc) were calculated for the predicted siRNAs. The higher values of these two melting temperatures indicate higher effectiveness of the siRNA species. Tm(Conc) values ranged from 71.7°C to 81.7°C for nucleocapsid phosphoprotein (Supplementary Table 6) and 66.4°C to 83.8°C for surface glycoprotein (Supplementary Table 7). Tm(Cp) values ranged from 72.1°C to 82.5°C for nucleocapsid phosphoprotein (Supplementary Table 6) and from 66.3°C to 85.2°C for surface glycoprotein (Supplementary Table 7).

### 3.7. Validation and selection of best 8 siRNAs

siRNAPred[53] checked the effectivity of the predicted siRNAs and values greater than 1 are considered highly effective. 2 siRNAs for nucleaocapsid phosphoprotein and 32 siRNAs for surface glycoprotein were found to be highly effective. Finally, based on all the other criteria, 8 siRNAs were selected as best predicted candidates against the nucleocapsid phosphoprotein and the surface glycoprotein genes of SARS CoV-2 (Table 2).

**Table 2.**
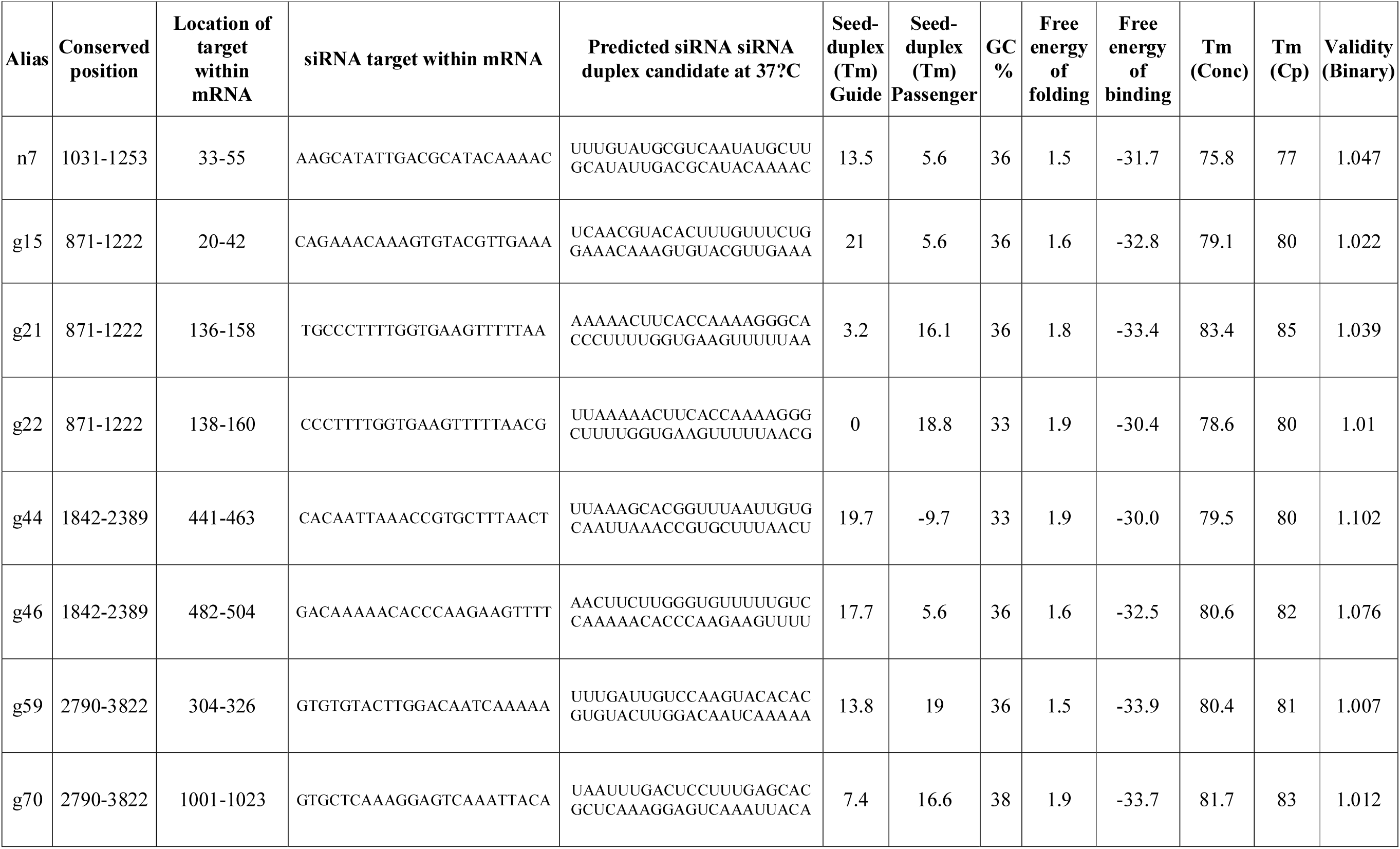
Best effective siRNA molecules with various parameters.

## 4. Discussion

COVID-19 is an emerging disease that lays bare the society we have created and its interdependent infrastructure with a surge in cases and deaths since its initial identification. Having no regard for geography, this pandemic has a global reach, and no continent is out of its clutches. Moreover, there is no vaccine available to prevent this disease and no RNAi based treatment is yet in practice or been proposed. So, the next generation medicine, siRNA might be effective in this case, hence it is the focus of our study.

Here, a total of 34 (15 nucleocapsid phosphoprotein and 19 surface glycoprotein) (Supplementary Table 2 & Supplementary table 3) conserved regions were identified among 139 strains of SARS-CoV-2 from around the world. Phylogenetic analysis revealed that a small number sequences form significant clades with a bootstrap value greater than 60%. (Fig 2, Supplementary Table 8). Conserved portions that are shorter than 21 nucleotides were omitted from further analysis. Conserved sequences were put to siDirect web server to identify possible targets and to generate corresponding siRNAs. siDirect performs the task in three distinct steps – highly functional siRNA selection, seed-dependent off-target effects reduction, near-perfect matched genes elimination. siRNA targets were found in 18 conserved regions, 5 nucleocapsid phosphoprotein (Supplementary Table 4) and 13 surface glycoprotein (Supplementary Table 5). U,R,A (Ui-Tei, Amarzguioui and Reynolds) rules (Table 1) were applied while predicting the siRNAs to obtain better results. siRNA bond formation with non-target sequences were eliminated by optimizing the melting temperature (Tm) below 21.5°C. The equation to calculate melting temperature (Tm) is below,

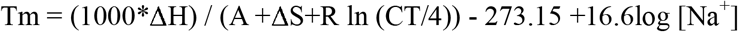

Here,

- The sum of the nearest neighbor enthalpy change is represented by ΔH (kcal*/* mol)
- The helix initiation constant (−10.8) is represented by A
- The sum of the nearest neighbor entropy change represented by ΔS
- The gas constant (1.987 cal/deg/mol) is represented by R
- The total molecular concentration (100 μM) of the strand is represented by CT and
- Concentration of Sodium, [Na^+^] was fixed at 100 mM

siRNA’s functionality is influenced by the GC-content and there is an inverse relationship of the GC-content with the function of siRNA. Usually a low GC content, approximately from 31.6 to 57.9%, is ideal for a siRNA to be effective [54]. Therefore, we calculated the GC content of the predicted siRNAs. Molecules that have GC content lower than 32% were not kept in the final selection. Here, GC content ranged from 10% to 43% for all the 78 predicted species. GC content of finally selected siRNAs are greater than or equal to 33% (Table 2).

Formation of secondary structure of siRNA may inhibit the RISC mediated cleavage of target. So, the prediction of prospective secondary structure and determination of free energy of corresponding folding is crucial. Here, guide strands of predicted siRNAs were subjected to RNA structure web server in order to predict possible folding structures and corresponding minimum free energies. At 37°C, finally selected siRNAs have free energy of folding greater than zero (Fig 3, Table 2), which suggests the predicted siRNAs are more accessible for efficient binding.

**Fig. 3.**
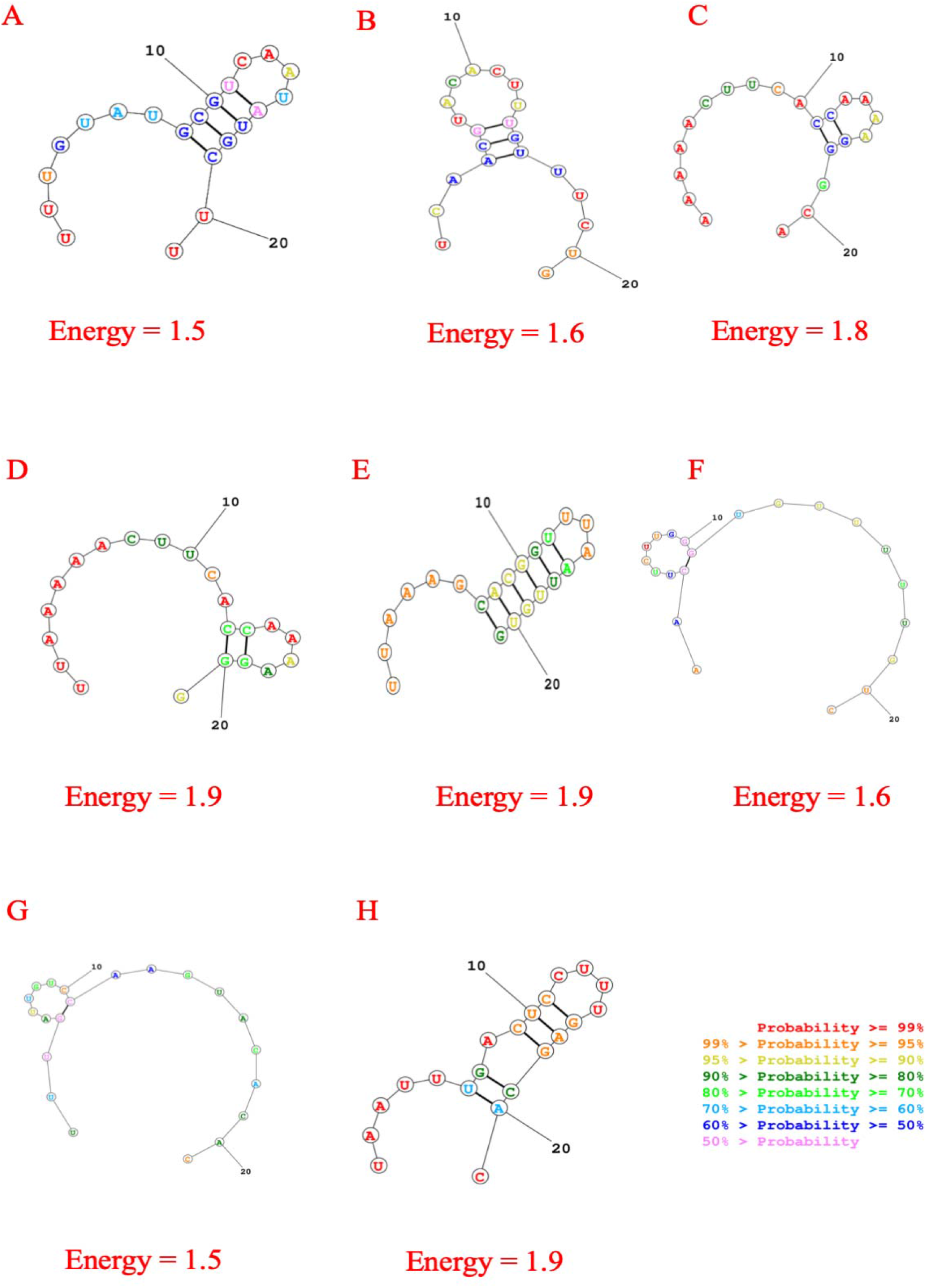
Secondary structures of best eight predicted siRNA with probable folding and lowest free energy for consensus sequence. The structures are for A. n7 B.g15 C. g21 D. g22 E. g44 F. g46 G. g59 H. g70 siRNAs.

DuplexFold[51] was used to determine the target and guide siRNA interaction and their corresponding binding energy. Lower binding energy indicate better interaction therefore better chance of target inhibition. The values of free energy of binding of all the 78 predicted siRNAs spanned from −36.6 to −21.6 (Supplementary Table 6 & Supplementary table 7). Finally selected siRNAs have free energy of binding equal or below −30.0 (Fig 4, Table 2), which suggests the predicted siRNAs are more interactive with their corresponding targets.

**Fig. 4.**
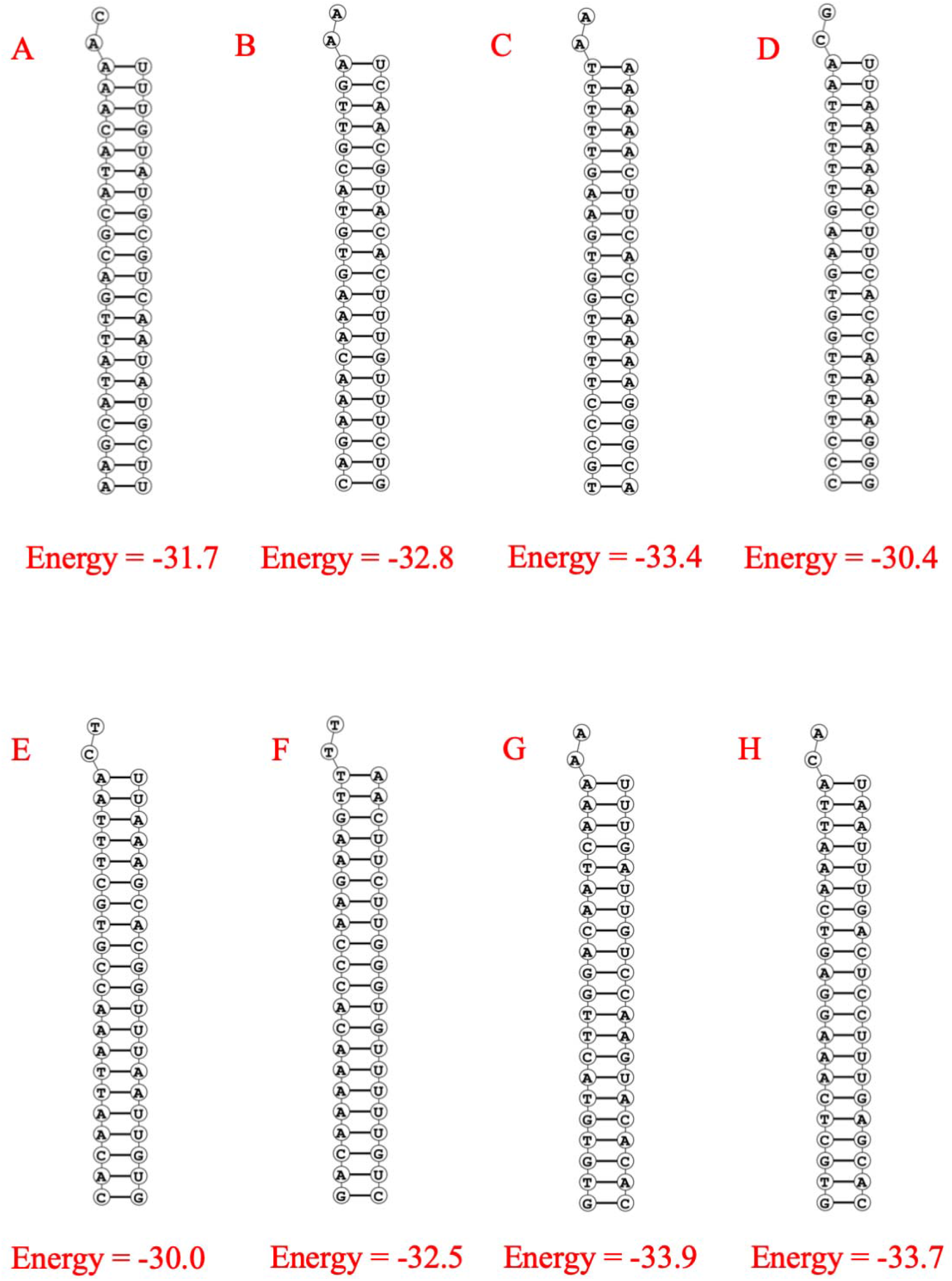
Structure of binding of siRNA (guide strand) and target RNA with corresponding predicted minimum free energy. The structures are for A. n7 B.g15 C. g21 D. g22 E. g44 F. g46 G. g59 H. g70 siRNAs and their corresponding targets.

The collective heat capacity, denoted as Cp, is plotted as a function of temperature and the melting temperature, denoted as Tm (Cp), was determined. The contribution of each molecules to the collective heat capacity was demonstrated using the inclusive heat capacity plot where melting temperature Tm (Conc), indicates the temperature at which the concentration of double-stranded molecules of becomes one-half of its maximum value. DINA Melt webserver [52] was used to obtain the melting temperatures. All the selected siRNAs have high Tm value (>75°C) (Table 2).

Lastly, siRNAPred[53] was used to determine the inhibition efficacy of the predicted siRNAs. siRNAPred uses Main21 dataset which consist of 2182 siRNAs (21mer) derived from a homogeneous experimental condition to predict the actual efficacy of 21mer siRNAs with high accuracy using the support vector machine based method. Here, siRNA candidates that have validity score greater than one were chosen for the final selection.

In this study, eight prospective siRNA molecules were proposed to be efficient at binding and cleaving specific mRNA targets of SARS-CoV-2 (Table 2). As the study contain a large array of 139 sequences of SARS-CoV-2 from around the world, the predicted therapeutic agent can be employed to large scale treatment of COVID-19 pandemic.

## 5. Conclusions

Computational methods can be employed to design and predict siRNA interaction against specific gene target thereby silencing its expression. In this research, eight siRNA molecules were predicted to be effective against nucleocapsid phosphoprotein and surface glycoprotein gene of 139 strains of SARS-CoV-2 virus using computational method considering all maximum parameters in prime conditions. In order to decelerate the COVID-19 pandemic and recover the affected individuals the development of siRNA therapeutic approaches could be a promising alternative to traditional vaccine designing.

## Author contribution statement

Literature Review: UFC, KIH, MUSS; Data Collection: UFC, MUSS; Data Analysis: UFC, MUSS; Figure: UFC, MKSS; Write-up: UFC, MUSS, KIH, MAB, MKSS, MAM;

## Supplement Table Captions

**Supplementary Table 1:** Accession number, Length and Location of the SARS-CoV-2 Stains Used in the Study.

**Supplementary Table 2:** Conserved sequence of nucleocapsid phosphoprotein gene.

**Supplementary Table 3:** Conserved sequence of surface glycoprotein gene.

**Supplementary Table 4:** Tm values of predicted siRNA (guide strand and passenger strand) against nucleocapsid phosphoprotein.

**Supplementary Table 5:** Tm values of predicted siRNA (guide strand and passenger strand) against surface glycoprotein.

**Supplementary Table 6:** Effective siRNAs against nucleocapsid phosphoprotein with GC%, free energy of folding and free energy of binding with target.

**Supplementary Table 7:** Effective siRNAs against surface glycoprotein with GC%, free energy of folding and free energy of binding with target.

**Supplementary Table 8:** Bootstrap values of significant clades in the radial phylogenetic tree.

