## Supplementary Table 1 for "A Computational Approach to Design Potential siRNA Molecules as a Prospective Tool for Silencing Nucleocapsid Phosphoprotein and Surface Glycoprotein Gene of SARS-CoV-2"

A Computational Approach to Design Potential siRNA Molecules as a Prospective Tool for Silencing Nucleocapsid Phosphoprotein and Surface Glycoprotein Gene of SARS-CoV-2

Umar Faruq Chowdhury^1^, Mohammad Umer Sharif Shohan^1^, Kazi Injamamul Hoque^1^ Mirza Ashikul Beg^1^, Mohammad Ali Moni^2^, Mohammad Kawsar Sharif Siam^3^

| **Table S1. Accession number, Length and Location of the SARS-CoV-2 Stains Used in the Study** | | | | | | | | |
| --- | --- | --- | --- | --- | --- | --- | --- | --- |
| **Accession** | **Length** | **Location** | **Accession** | **Length** | **Location** | **Accession** | **Length** | **Location** |
| MT233526 | 29847 | USA | MT233522 | 29782 | Spain | MT135042 | 29903 | China |
| MT246449 | 29828 | USA | MT233523 | 29782 | Spain | MT135043 | 29903 | China |
| MT246450 | 29872 | USA | MT226610 | 29899 | China | MT135044 | 29903 | China |
| MT246451 | 29868 | USA | MT192772 | 29891 | Viet Nam | MT126808 | 29876 | Brazil |
| MT246452 | 29888 | USA | MT192765 | 29829 | USA | MT123292 | 29923 | China |
| MT246453 | 29823 | USA | MT192759 | 29862 | Taiwan | MT123290 | 29891 | China |
| MT246454 | 29892 | USA | MT192773 | 29890 | Viet Nam | MT123291 | 29882 | China |
| MT246455 | 29857 | USA | MT188340 | 29845 | USA | MT123293 | 29871 | China |
| MT246456 | 29846 | USA | MT188339 | 29783 | USA | MT118835 | 29882 | USA |
| MT246457 | 29843 | USA | MT188341 | 29835 | USA | MT106053 | 29882 | USA |
| MT246458 | 29765 | USA | MT184908 | 29880 | USA | MT106052 | 29882 | USA |
| MT246459 | 29909 | USA | MT184909 | 29882 | USA | MT106054 | 29882 | USA |
| MT246460 | 29917 | USA | MT184910 | 29882 | USA | MT093631 | 29860 | China |
| MT246461 | 29877 | USA | MT184907 | 29882 | USA | MT093571 | 29886 | Sweden |
| MT246462 | 29903 | USA | MT184911 | 29882 | USA | MT072688 | 29811 | Nepal |
| MT246463 | 29782 | USA | MT184913 | 29882 | USA | MT066175 | 29870 | Taiwan |
| MT246464 | 29818 | USA | MT184912 | 29882 | USA | MT066176 | 29870 | Taiwan |
| MT246466 | 29897 | USA | MT163719 | 29903 | USA | MT049951 | 29903 | China |
| MT246467 | 29899 | USA | MT163716 | 29903 | USA | MT044257 | 29882 | USA |
| MT246468 | 29843 | USA | MT163717 | 29897 | USA | MT044258 | 29858 | USA |
| MT246469 | 29869 | USA | MT163718 | 29903 | USA | MT039890 | 29903 | South Korea |
| MT246470 | 29860 | USA | MT159705 | 29882 | USA | MT039887 | 29879 | USA |
| MT246471 | 29871 | USA | MT159722 | 29882 | USA | MT039873 | 29833 | China |
| MT246472 | 29810 | USA | MT159710 | 29882 | USA | MT039888 | 29882 | USA |
| MT246473 | 29825 | USA | MT159715 | 29882 | USA | MT027063 | 29882 | USA |
| MT246474 | 29884 | USA | MT159716 | 29867 | USA | MT027064 | 29882 | USA |
| MT246475 | 29877 | USA | MT159711 | 29882 | USA | MT027062 | 29882 | USA |
| MT246476 | 29863 | USA | MT159712 | 29882 | USA | MT020881 | 29882 | USA |
| MT246477 | 29886 | USA | MT159717 | 29882 | USA | MT019533 | 29883 | China |
| MT246478 | 29889 | USA | MT159719 | 29882 | USA | MT019531 | 29899 | China |
| MT246479 | 29828 | USA | MT159720 | 29882 | USA | MT019532 | 29890 | China |
| MT246480 | 29904 | USA | MT159709 | 29882 | USA | MT019530 | 29889 | China |
| MT246481 | 29873 | USA | MT121215 | 29945 | China | MT019529 | 29899 | China |
| MT246482 | 29844 | USA | MT159718 | 29882 | USA | MT020880 | 29882 | USA |
| MT246484 | 29870 | USA | MT159721 | 29882 | USA | MT007544 | 29893 | Australia |
| MT246485 | 29775 | USA | MT159708 | 29882 | USA | MN996527 | 29825 | China |
| MT246486 | 29867 | USA | MT159714 | 29882 | USA | MN996528 | 29891 | China |

| **Table S1. Accession number, Length and Location of the SARS-CoV-2 Stains Used in the Study (Continued)** | | | | | | | | |
| --- | --- | --- | --- | --- | --- | --- | --- | --- |
| **Accession** | **Length** | **Location** | **Accession** | **Length** | **Location** | **Accession** | **Length** | **Location** |
| MT246487 | 29883 | USA | MT159707 | 29882 | USA | MN996529 | 29852 | China |
| MT246488 | 29842 | USA | MT066156 | 29867 | Italy | MN996530 | 29854 | China |
| MT246489 | 29864 | USA | MT159706 | 29882 | USA | MN996531 | 29857 | China |
| MT246490 | 29850 | USA | MT159713 | 29882 | USA | MN988669 | 29881 | China |
| MT246667 | 29867 | USA | MT012098 | 29854 | India | MN994467 | 29882 | USA |
| MT240479 | 29836 | Pakistan | MT050493 | 29851 | India | MN994468 | 29883 | USA |
| MT233519 | 29782 | Spain | MT152824 | 29878 | USA | MN997409 | 29882 | USA |
| MN988713 | 29882 | USA | MT135041 | 29903 | China | MN988668 | 29881 | China |
| MN975262 | 29891 | China | MN938384 | 29838 | China | MN985325 | 29882 | USA |
| MN908947 | 29903 | China |  |  |  |  |  |  |
