## Supplementary Table 2 for "A Computational Approach to Design Potential siRNA Molecules as a Prospective Tool for Silencing Nucleocapsid Phosphoprotein and Surface Glycoprotein Gene of SARS-CoV-2"

A Computational Approach to Design Potential siRNA Molecules as a Prospective Tool for Silencing Nucleocapsid Phosphoprotein and Surface Glycoprotein Gene of SARS-CoV-2

Umar Faruq Chowdhury^1^, Mohammad Umer Sharif Shohan^1^, Kazi Injamamul Hoque^1^ Mirza Ashikul Beg^1^, Mohammad Ali Moni^2^, Mohammad Kawsar Sharif Siam^3^

| **Table S2. Conserved sequence of nucleocapsid phosphoprotein gene** | |
| --- | --- |
| **Position** | **Sequence** |
| 1-104 | ATGTCTGATAATGGACCCCAAAATCAGCGAAATGCACCCCGCATTACGTTTGGTGGACCC TCAGATTCAACTGGCAGTAACCAGAATGGAGAACGCAGTGGGGC |
| 106-135 | CGATCAAAACAACGTCGGCCCCAAGGTTTA |
| 137-383 | CCAATAATACTGCGTCTTGGTTCACCGCTCTCACTCAACATGGCAAGGAAGACCTTAAAT TCCCTCGAGGACAAGGCGTTCCAATTAACACCAATAGCAGTCCAGATGACCAAATTGGCT ACTACCGAAGAGCTACCAGACGAATTCGTGGTGGTGACGGTAAAATGAAAGATCTCAGTC CAAGATGGTATTTCTACTACCTAGGAACTGGGCCAGAAGCTGGACTTCCCTATGGTGCTA ACAAAGA |
| 385-518 | GGCATCATATGGGTTGCAACTGAGGGAGCCTTGAATACACCAAAAGATCACATTGGCACC CGCAATCCTGCTAACAATGCTGCAATCGTGCTACAACTTCCTCAAGGAACAACATTGCCA AAAGGCTTCTACGC |
| 520-552 | GAAGGGAGCAGAGGCGGCAGTCAAGCCTCTTCT |
| 554-580 | GTTCCTCATCACGTAGTCGCAACAGTT |
| 611-642 | GAACTTCTCCTGCTAGAATGGCTGGCAATGGC |
| 644-821 | GTGATGCTGCTCTTGCTTTGCTGCTGCTTGACAGATTGAACCAGCTTGAGAGCAAAATGT CTGGTAAAGGCCAACAACAACAAGGCCAAACTGTCACTAAGAAATCTGCTGCTGAGGCTT CTAAGAAGCCTCGGCAAAAACGTACTGCCACTAAAGCATACAATGTAACACAAGCTTT |
| 823-956 | GGCAGACGTGGTCCAGAACAAACCCAAGGAAATTTTGGGGACCAGGAACTAATCAGACAA GGAACTGATTACAAACATTGGCCGCAAATTGCACAATTTGCCCCCAGCGCTTCAGCGTTC TTCGGAATGTCGCG |
| 958-980 | ATTGGCATGGAAGTCACACCTTC |
| 982-1027 | GGAACGTGGTTGACCTACACAGGTGCCATCAAATTGGATGACAAAG |
| 1029 | T |
| 1031-1253 | CAAATTTCAAAGATCAAGTCATTTTGCTGAATAAGCATATTGACGCATACAAAACATTCC CACCAACAGAGCCTAAAAAGGACAAAAAGAAGAAGGCTGATGAAACTCAAGCCTTACCGC AGAGACAGAAGAAACAGCAAACTGTGACTCTTCTTCCTGCTGCAGATTTGGATGATTTCT CCAAACAATTGCAACAATCCATGAGCAGTGCTGACTCAACTCA |
| 1255-1260 | GCCTAA |
