## Supplementary Table 3 for "A Computational Approach to Design Potential siRNA Molecules as a Prospective Tool for Silencing Nucleocapsid Phosphoprotein and Surface Glycoprotein Gene of SARS-CoV-2"

A Computational Approach to Design Potential siRNA Molecules as a Prospective Tool for Silencing Nucleocapsid Phosphoprotein and Surface Glycoprotein Gene of SARS-CoV-2

Umar Faruq Chowdhury^1^, Mohammad Umer Sharif Shohan^1^, Kazi Injamamul Hoque^1^ Mirza Ashikul Beg^1^, Mohammad Ali Moni^2^, Mohammad Kawsar Sharif Siam^3^

| **Table S3. Conserved sequence of surface glycoprotein gene** | |
| --- | --- |
| **Position** | **Sequence** |
| 1-12 | ATGTTTGTTTTT |
| 14-81 | TTGTTTTATTGCCACTAGTCTCTAGTCAGTGTGTTAATCTTACAACCAGAACTCAATTAC CCCCTGCA |
| 83-144 | ACACTAATTCTTTCACACGTGGTGTTTATTACCCTGACAAAGTTTTCAGATCCTCAGTTT TA |
| 146-221 | ATTCAACTCAGGACTTGTTCTTACCTTTCTTTTCCAATGTTACTTGGTTCCATGCTATAC ATGTCTCTGGGACCAA |
| 223-470 | GGTACTAAGAGGTTTGATAACCCTGTCCTACCATTTAATGATGGTGTTTATTTTGCTTCC ACTGAGAAGTCTAACATAATAAGAGGCTGGATTTTTGGTACTACTTTAGATTCGAAGACC CAGTCCCTACTTATTGTTAATAACGCTACTAATGTTGTTATTAAAGTCTGTGAATTTCAA TTTTGTAATGATCCATTTTTGGGTGTTTATTACCACAAAAACAACAAAAGTTGGATGGAA AGTGAGTT |
| 472-541 | AGAGTTTATTCTAGTGCGAATAATTGCACTTTTGAATATGTCTCTCAGCCTTTTCTTATG GACCTTGAAG |
| 543-661 | AAAACAGGGTAATTTCAAAAATCTTAGGGAATTTGTGTTTAAGAATATTGATGGTTATTT TAAAATATATTCTAAGCACACGCCTATTAATTTAGTGCGTGATCTCCCTCAGGGTTTTT |
| 663-740 | GGCTTTAGAACCATTGGTAGATTTGCCAATAGGTATTAACATCACTAGGTTTCAAACTTT ACTTGCTTTACATAGAAG |
| 742-869 | TATTTGACTCCTGGTGATTCTTCTTCAGGTTGGACAGCTGGTGCTGCAGCTTATTATGTG GGTTATCTTCAACCTAGGACTTTTCTATTAAAATATAATGAAAATGGAACCATTACAGAT GCTGTAGA |
| 871-1222 | TGTGCACTTGACCCTCTCTCAGAAACAAAGTGTACGTTGAAATCCTTCACTGTAGAAAAA GGAATCTATCAAACTTCTAACTTTAGAGTCCAACCAACAGAATCTATTGTTAGATTTCCT AATATTACAAACTTGTGCCCTTTTGGTGAAGTTTTTAACGCCACCAGATTTGCATCTGTT TATGCTTGGAACAGGAAGAGAATCAGCAACTGTGTTGCTGATTATTCTGTCCTATATAAT TCCGCATCATTTTCCACTTTTAAGTGTTATGGAGTGTCTCCTACTAAATTAAATGATCTC TGCTTTACTAATGTCTATGCAGATTCATTTGTAATTAGAGGTGATGAAGTCA |
| 1224-1421 | ACAAATCGCTCCAGGGCAAACTGGAAAGATTGCTGATTATAATTATAAATTACCAGATGA TTTTACAGGCTGCGTTATAGCTTGGAATTCTAACAATCTTGATTCTAAGGTTGGTGGTAA TTATAATTACCTGTATAGATTGTTTAGGAAGTCTAATCTCAAACCTTTTGAGAGAGATAT TTCAACTGAAATCTATCA |
| 1423-1425 | GCC |
| 1427-1622 | GTAGCACACCTTGTAATGGTGTTGAAGGTTTTAATTGTTACTTTCCTTTACAATCATATG GTTTCCAACCCACTAATGGTGTTGGTTACCAACCATACAGAGTAGTAGTACTTTCTTTTG AACTTCTACATGCACCAGCAACTGTTTGTGGACCTAAAAAGTCTACTAATTTGGTTAAAA ACAAATGTGTCAATTT |

| **Table S3. Conserved sequence of surface glycoprotein gene (Continued)** | |
| --- | --- |
| **Position** | **Sequence** |
| 1624-1840 | AACTTCAATGGTTTAACAGGCACAGGTGTTCTTACTGAGTCTAACAAAAAGTTTCTGCCT TTCCAACAATTTGGCAGAGACATTGCTGACACTACTGATGCTGTCCGTGATCCACAGACA CTTGAGATTCTTGACATTACACCATGTTCTTTTGGTGGTGTCAGTGTTATAACACCAGGA ACAAATACTTCTAACCAGGTTGCTGTTCTTTATCAGG |
| 1842-2389 | TGTTAACTGCACAGAAGTCCCTGTTGCTATTCATGCAGATCAACTTACTCCTACTTGGCG TGTTTATTCTACAGGTTCTAATGTTTTTCAAACACGTGCAGGCTGTTTAATAGGGGCTGA ACATGTCAACAACTCATATGAGTGTGACATACCCATTGGTGCAGGTATATGCGCTAGTTA TCAGACTCAGACTAATTCTCCTCGGCGGGCACGTAGTGTAGCTAGTCAATCCATCATTGC CTACACTATGTCACTTGGTGCAGAAAATTCAGTTGCTTACTCTAATAACTCTATTGCCAT ACCCACAAATTTTACTATTAGTGTTACCACAGAAATTCTACCAGTGTCTATGACCAAGAC ATCAGTAGATTGTACAATGTACATTTGTGGTGATTCAACTGAATGCAGCAATCTTTTGTT GCAATATGGCAGTTTTTGTACACAATTAAACCGTGCTTTAACTGGAATAGCTGTTGAACA AGACAAAAACACCCAAGAAGTTTTTGCACAAGTCAAACAAATTTACAAAACACCACCAAT TAAAGATT |
| 2391-2471 | TGGTGGTTTTAATTTTTCACAAATATTACCAGATCCATCAAAACCAAGCAAGAGGTCATT TATTGAAGATCTACTTTTCAA |
| 2473-2762 | AAAGTGACACTTGCAGATGCTGGCTTCATCAAACAATATGGTGATTGCCTTGGTGATATT GCTGCTAGAGACCTCATTTGTGCACAAAAGTTTAACGGCCTTACTGTTTTGCCACCTTTG CTCACAGATGAAATGATTGCTCAATACACTTCTGCACTGTTAGCGGGTACAATCACTTCT GGTTGGACCTTTGGTGCAGGTGCTGCATTACAAATACCATTTGCTATGCAAATGGCTTAT AGGTTTAATGGTATTGGAGTTACACAGAATGTTCTCTATGAGAACCAAAA |
| 2764-2788 | TTGATTGCCAACCAATTTAATAGTG |
| 2790-3822 | TATTGGCAAAATTCAAGACTCACTTTCTTCCACAGCAAGTGCACTTGGAAAACTTCAAGA TGTGGTCAACCAAAATGCACAAGCTTTAAACACGCTTGTTAAACAACTTAGCTCCAATTT TGGTGCAATTTCAAGTGTTTTAAATGATATCCTTTCACGTCTTGACAAAGTTGAGGCTGA AGTGCAAATTGATAGGTTGATCACAGGCAGACTTCAAAGTTTGCAGACATATGTGACTCA ACAATTAATTAGAGCTGCAGAAATCAGAGCTTCTGCTAATCTTGCTGCTACTAAAATGTC AGAGTGTGTACTTGGACAATCAAAAAGAGTTGATTTTTGTGGAAAGGGCTATCATCTTAT GTCCTTCCCTCAGTCAGCACCTCATGGTGTAGTCTTCTTGCATGTGACTTATGTCCCTGC ACAAGAAAAGAACTTCACAACTGCTCCTGCCATTTGTCATGATGGAAAAGCACACTTTCC TCGTGAAGGTGTCTTTGTTTCAAATGGCACACACTGGTTTGTAACACAAAGGAATTTTTA TGAACCACAAATCATTACTACAGACAACACATTTGTGTCTGGTAACTGTGATGTTGTAAT AGGAATTGTCAACAACACAGTTTATGATCCTTTGCAACCTGAATTAGACTCATTCAAGGA GGAGTTAGATAAATATTTTAAGAATCATACATCACCAGATGTTGATTTAGGTGACATCTC TGGCATTAATGCTTCAGTTGTAAACATTCAAAAAGAAATTGACCGCCTCAATGAGGTTGC CAAGAATTTAAATGAATCTCTCATCGATCTCCAAGAACTTGGAAAGTATGAGCAGTATAT AAAATGGCCATGGTACATTTGGCTAGGTTTTATAGCTGGCTTGATTGCCATAGTAATGGT GACAATTATGCTTTGCTGTATGACCAGTTGCTGTAGTTGTCTCAAGGGCTGTTGTTCTTG TGGATCCTGCTGCAAATTTGATGAAGACGACTCTGAGCCAGTGCTCAAAGGAGTCAAATT ACATTACACATAA |
