## Supplementary Table 4 for "A Computational Approach to Design Potential siRNA Molecules as a Prospective Tool for Silencing Nucleocapsid Phosphoprotein and Surface Glycoprotein Gene of SARS-CoV-2"

A Computational Approach to Design Potential siRNA Molecules as a Prospective Tool for Silencing Nucleocapsid Phosphoprotein and Surface Glycoprotein Gene of SARS-CoV-2

Umar Faruq Chowdhury^1^, Mohammad Umer Sharif Shohan^1^, Kazi Injamamul Hoque^1^ Mirza Ashikul Beg^1^, Mohammad Ali Moni^2^, Mohammad Kawsar Sharif Siam^3^

| **Table S4. Tm values of predicted siRNA (guide strand and passenger strand) against nucleocapsid phosphoprotein.** | | | |
| --- | --- | --- | --- |
| **Alias** | **Predicted siRNA siRNA  duplex candidate at 37◦C; RNA oligo sequences  21nt guide (5' →3' ) 21nt passenger (5' →3' )** | **Seed-duplex Tm, ◦C** | |
|  |  | **Guide** | **Passenger** |
| n1 | AGUAGAAAUACCAUCUUGGAC CCAAGAUGGUAUUUCUACUAC | 14.6 | 18.1 |
| n2 | UCUUUUGGUGUAUUCAAGGCU CCUUGAAUACACCAAAAGAUC | 16.1 | 12 |
| n3 | UUUCUUAGUGACAGUUUGGCC CCAAACUGUCACUAAGAAAUC | 11.7 | 16.7 |
| n4 | ACAUUGUAUGCUUUAGUGGCA CCACUAAAGCAUACAAUGUAA | 13.5 | 11.8 |
| n5 | AAUUUCCUUGGGUUUGUUCUG GAACAAACCCAAGGAAAUUUU | 18.7 | 13.3 |
| n6 | AAUUUGCGGCCAAUGUUUGUA CAAACAUUGGCCGCAAAUUGC | 20.6 | 5.3 |
| n7 | UUUGUAUGCGUCAAUAUGCUU GCAUAUUGACGCAUACAAAAC | 13.5 | 5.6 |
| n8 | UUCUUUUUGUCCUUUUUAGGC CUAAAAAGGACAAAAAGAAGA | 5.5 | -3.8 |
