## Supplementary Table 5 for "A Computational Approach to Design Potential siRNA Molecules as a Prospective Tool for Silencing Nucleocapsid Phosphoprotein and Surface Glycoprotein Gene of SARS-CoV-2"

A Computational Approach to Design Potential siRNA Molecules as a Prospective Tool for Silencing Nucleocapsid Phosphoprotein and Surface Glycoprotein Gene of SARS-CoV-2

Umar Faruq Chowdhury^1^, Mohammad Umer Sharif Shohan^1^, Kazi Injamamul Hoque^1^ Mirza Ashikul Beg^1^, Mohammad Ali Moni^2^, Mohammad Kawsar Sharif Siam^3^

| **Table S5. Tm values of predicted siRNA (guide strand and passenger strand) against surface glycoprotein.** | | | |
| --- | --- | --- | --- |
| **Alias** | **Predicted siRNA siRNA  duplex candidate at 37◦C; RNA oligo sequences  21nt guide (5' →3' ) 21nt passenger (5' →3' )** | **Seed-duplex Tm, ◦C** | |
|  |  | **Guide** | **Passenger** |
| g1 | UAACAUUGGAAAAGAAAGGUA CCUUUCUUUUCCAAUGUUACU | 12.1 | 10.3 |
| g2 | UAAAGUAGUACCAAAAAUCCA GAUUUUUGGUACUACUUUAGA | 9.8 | -3.3 |
| g3 | UUAAUAACAACAUUAGUAGCG CUACUAAUGUUGUUAUUAAAG | 1.4 | 6.3 |
| g4 | UAAUAAACACCCAAAAAUGGA CAUUUUUGGGUGUUUAUUACC | -0.3 | -3.3 |
| g5 | UUUUUGUGGUAAUAAACACCC GUGUUUAUUACCACAAAAACA | 12.2 | 6.9 |
| g6 | UCAAAAGUGCAAUUAUUCGCA CGAAUAAUUGCACUUUUGAAU | 10.3 | 1.8 |
| g7 | UUCAAAAGUGCAAUUAUUCGC GAAUAAUUGCACUUUUGAAUA | 12.2 | -10.3 |
| g8 | UUCUUAAACACAAAUUCCCUA GGGAAUUUGUGUUUAAGAAUA | 7.1 | 14.2 |
| g9 | UAUUCUUAAACACAAAUUCCC GAAUUUGUGUUUAAGAAUAUU | 6.9 | 5.3 |
| g10 | AAAAUAACCAUCAAUAUUCUU GAAUAUUGAUGGUUAUUUUAA | -0.3 | -1.8 |
| g11 | AUAUUUUAAAAUAACCAUCAA GAUGGUUAUUUUAAAAUAUAU | -7.5 | 20 |
| g12 | UAUUUUAAUAGAAAAGUCCUA GGACUUUUCUAUUAAAAUAUA | -9.7 | 13.3 |
| g13 | UUAUAUUUUAAUAGAAAAGUC CUUUUCUAUUAAAAUAUAAUG | -8 | 7.1 |
| g14 | UUUCAUUAUAUUUUAAUAGAA CUAUUAAAAUAUAAUGAAAAU | 8.9 | -7.5 |
| g15 | UCAACGUACACUUUGUUUCUG GAAACAAAGUGUACGUUGAAA | 21 | 5.6 |
| g16 | UUUGAUAGAUUCCUUUUUCUA GAAAAAGGAAUCUAUCAAACU | 13.4 | 9.5 |
| g17 | UAGAAGUUUGAUAGAUUCCUU GGAAUCUAUCAAACUUCUAAC | 17.7 | 16 |
| g18 | UUAGAAGUUUGAUAGAUUCCU GAAUCUAUCAAACUUCUAACU | 18.9 | 6.6 |
| g19 | AAAGUUAGAAGUUUGAUAGAU CUAUCAAACUUCUAACUUUAG | 9.8 | 8.9 |
| g20 | UAAUAUUAGGAAAUCUAACAA GUUAGAUUUCCUAAUAUUACA | -8 | 6.9 |

| **Table S5. Tm values of predicted siRNA (guide strand and passenger strand) against surface glycoprotein. (continued)** | | | |
| --- | --- | --- | --- |
| **Alias** | **Predicted siRNA siRNA  duplex candidate at 37◦C; RNA oligo sequences  21nt guide (5' →3' ) 21nt passenger (5' →3' )** | **Seed-duplex Tm, ◦C** | |
|  |  | **Guide** | **Passenger** |
| g21 | AAAAACUUCACCAAAAGGGCA CCCUUUUGGUGAAGUUUUUAA | 3.2 | 16.1 |
| g22 | UUAAAAACUUCACCAAAAGGG CUUUUGGUGAAGUUUUUAACG | 0 | 18.8 |
| g23 | UAAACAGAUGCAAAUCUGGUG CCAGAUUUGCAUCUGUUUAUG | 19.2 | 12 |
| g24 | UAUAUAGGACAGAAUAAUCAG GAUUAUUCUGUCCUAUAUAAU | 12.2 | 1.8 |
| g25 | UAAAAGUGGAAAAUGAUGCGG GCAUCAUUUUCCACUUUUAAG | 10.3 | 13.6 |
| g26 | UUAAAAGUGGAAAAUGAUGCG CAUCAUUUUCCACUUUUAAGU | 4.9 | 7.2 |
| g27 | UAAUUACAAAUGAAUCUGCAU GCAGAUUCAUUUGUAAUUAGA | 6.9 | 20.4 |
| g28 | UAUAAUCAGCAAUCUUUCCAG GGAAAGAUUGCUGAUUAUAAU | 8.7 | 14.8 |
| g29 | UUAUAAUCAGCAAUCUUUCCA GAAAGAUUGCUGAUUAUAAUU | 3.5 | 5.3 |
| g30 | UAGAAUUCCAAGCUAUAACGC GUUAUAGCUUGGAAUUCUAAC | 14.8 | 14.9 |
| g31 | UUAGAAUCAAGAUUGUUAGAA CUAACAAUCUUGAUUCUAAGG | 16 | 6.9 |
| g32 | UAAUUACCACCAACCUUAGAA CUAAGGUUGGUGGUAAUUAUA | 13.9 | 18.6 |
| g33 | UUUUUAGGUCCACAAACAGUU CUGUUUGUGGACCUAAAAAGU | 11 | 19.3 |
| g34 | AAUUAGUAGACUUUUUAGGUC CCUAAAAAGUCUACUAAUUUG | 6.3 | -3.8 |
| g35 | AAAUUAGUAGACUUUUUAGGU CUAAAAAGUCUACUAAUUUGG | 4.6 | -3.8 |
| g36 | UUUAACCAAAUUAGUAGACUU GUCUACUAAUUUGGUUAAAAA | 20 | 20.1 |
| g37 | UUUUUAACCAAAUUAGUAGAC CUACUAAUUUGGUUAAAAACA | 0 | 6.3 |
| g38 | UUUUGUUAGACUCAGUAAGAA CUUACUGAGUCUAACAAAAAG | 7.2 | 20.3 |
| g39 | AUAGUGUAGGCAAUGAUGGAU CCAUCAUUGCCUACACUAUGU | 20.2 | 13.6 |

| **Table S5. Tm values of predicted siRNA (guide strand and passenger strand) against surface glycoprotein. (continued)** | | | |
| --- | --- | --- | --- |
| **Alias** | **Predicted siRNA siRNA  duplex candidate at 37◦C; RNA oligo sequences  21nt guide (5' →3' ) 21nt passenger (5' →3' )** | **Seed-duplex Tm, ◦C** | |
|  |  | **Guide** | **Passenger** |
| g40 | UAUUGCAACAAAAGAUUGCUG GCAAUCUUUUGUUGCAAUAUG | 20 | 12 |
| g41 | ACAAAAACUGCCAUAUUGCAA GCAAUAUGGCAGUUUUUGUAC | 5.6 | 5.6 |
| g42 | UAAUUGUGUACAAAAACUGCC CAGUUUUUGUACACAAUUAAA | 12.1 | 3.2 |
| g43 | UUUAAUUGUGUACAAAAACUG GUUUUUGUACACAAUUAAACC | -1.4 | 5.6 |
| g44 | UUAAAGCACGGUUUAAUUGUG CAAUUAAACCGUGCUUUAACU | 19.7 | -9.7 |
| g45 | UUUUUGUCUUGUUCAACAGCU CUGUUGAACAAGACAAAAACA | 14.9 | 20.5 |
| g46 | AACUUCUUGGGUGUUUUUGUC CAAAAACACCCAAGAAGUUUU | 17.7 | 5.6 |
| g47 | UGUUUUGUAAAUUUGUUUGAC CAAACAAAUUUACAAAACACC | 13.3 | 5.6 |
| g48 | UUUAAUUGGUGGUGUUUUGUA CAAAACACCACCAAUUAAAGA | -1.4 | 13.3 |
| g49 | UUUGUGAAAAAUUAAAACCAC GGUUUUAAUUUUUCACAAAUA | 20.5 | 0 |
| g50 | UAGAUCUUCAAUAAAUGACCU GUCAUUUAUUGAAGAUCUACU | 19.1 | 8.9 |
| g51 | AAACCUAUAAGCCAUUUGCAU GCAAAUGGCUUAUAGGUUUAA | 18.5 | 17.7 |
| g52 | UCAUAGAGAACAUUCUGUGUA CACAGAAUGUUCUCUAUGAGA | 17.8 | 19.2 |
| g53 | UAAAGCUUGUGCAUUUUGGUU CCAAAAUGCACAAGCUUUAAA | 17 | 4.2 |
| g54 | UUAAAGCUUGUGCAUUUUGGU CAAAAUGCACAAGCUUUAAAC | 18.3 | 14 |
| g55 | UUAAAACACUUGAAAUUGCAC GCAAUUUCAAGUGUUUUAAAU | 7.2 | 7.4 |
| g56 | UUUAAAACACUUGAAAUUGCA CAAUUUCAAGUGUUUUAAAUG | 0 | 7.4 |

| **Table S5. Tm values of predicted siRNA (guide strand and passenger strand) against surface glycoprotein. (continued)** | | | |
| --- | --- | --- | --- |
| **Alias** | **Predicted siRNA siRNA  duplex candidate at 37◦C; RNA oligo sequences  21nt guide (5' →3' ) 21nt passenger (5' →3' )** | **Seed-duplex Tm, ◦C** | |
|  |  | **Guide** | **Passenger** |
| g57 | AUAUCAUUUAAAACACUUGAA CAAGUGUUUUAAAUGAUAUCC | 8.7 | 17.8 |
| g58 | UUUGUCAAGACGUGAAAGGAU CCUUUCACGUCUUGACAAAGU | 20.5 | 19.2 |
| g59 | UUUGAUUGUCCAAGUACACAC GUGUACUUGGACAAUCAAAAA | 13.8 | 19 |
| g60 | UUUUCCAUCAUGACAAAUGGC CAUUUGUCAUGAUGGAAAAGC | 20.1 | 14.8 |
| g61 | AAAAAUUCCUUUGUGUUACAA GUAACACAAAGGAAUUUUUAU | 0.4 | 20.4 |
| g62 | UGAAUGAGUCUAAUUCAGGUU CCUGAAUUAGACUCAUUCAAG | 20.4 | 12 |
| g63 | UCUAACUCCUCCUUGAAUGAG CAUUCAAGGAGGAGUUAGAUA | 18.9 | 12 |
| g64 | AAAAUAUUUAUCUAACUCCUC GGAGUUAGAUAAAUAUUUUAA | -10.3 | 18.9 |
| g65 | UAAAAUAUUUAUCUAACUCCU GAGUUAGAUAAAUAUUUUAAG | -10.3 | 18.9 |
| g66 | UUUCUUUUUGAAUGUUUACAA GUAAACAUUCAAAAAGAAAUU | 5.5 | 6.9 |
| g67 | AUACUUUCCAAGUUCUUGGAG CCAAGAACUUGGAAAGUAUGA | 14.6 | 19.2 |
| g68 | AUAUACUGCUCAUACUUUCCA GAAAGUAUGAGCAGUAUAUAA | 13.1 | 4.6 |
| g69 | AAAACCUAGCCAAAUGUACCA GUACAUUUGGCUAGGUUUUAU | 18.6 | 6.9 |
| g70 | UAAUUUGACUCCUUUGAGCAC GCUCAAAGGAGUCAAAUUACA | 7.4 | 16.6 |
