## Supplementary Table 6 for "A Computational Approach to Design Potential siRNA Molecules as a Prospective Tool for Silencing Nucleocapsid Phosphoprotein and Surface Glycoprotein Gene of SARS-CoV-2"

A Computational Approach to Design Potential siRNA Molecules as a Prospective Tool for Silencing Nucleocapsid Phosphoprotein and Surface Glycoprotein Gene of SARS-CoV-2

Umar Faruq Chowdhury^1^, Mohammad Umer Sharif Shohan^1^, Kazi Injamamul Hoque^1^ Mirza Ashikul Beg^1^, Mohammad Ali Moni^2^, Mohammad Kawsar Sharif Siam^3^

| **Table S6. Effective siRNAs against nucleocapsid phosphoprotein with GC%, free energy of folding and free energy of binding with target.** | | | | | | | | | |
| --- | --- | --- | --- | --- | --- | --- | --- | --- | --- |
| **Alias** | **Conserved  position** | **Location of target  within mRNA** | **siRNA target within mRNA** | **GC %** | **Free energy of folding** | **Free energy  of binding** | **Tm (conc)** | **Tm (Cp)** | **Validity  (binary)** |
| n1 | 137-383 | 178-200 | GTCCAAGATGGTATTTCTACTAC | 38 | 1.7 | -34.5 | 73.4 | 72.1 | 0.985 |
| n2 | 385-518 | 27-49 | AGCCTTGAATACACCAAAAGATC | 38 | 1.8 | -34.2 | 74.6 | 75.5 | 0.986 |
| n3 | 644-821 | 84-106 | GGCCAAACTGTCACTAAGAAATC | 40 | 1.8 | -35.8 | 75.4 | 76.5 | 0.959 |
| n4 | 644-821 | 146-168 | TGCCACTAAAGCATACAATGTAA | 36 | 1.7 | -34.5 | 75.2 | 75.3 | 0.986 |
| n5 | 823-956 | 14-36 | CAGAACAAACCCAAGGAAATTTT | 36 | 1.9 | -32.5 | 78.2 | 78.1 | 0.934 |
| n6 | 823-956 | 70-92 | TACAAACATTGGCCGCAAATTGC | 43 | 1.4 | -33.1 | 81.7 | 82.5 | 0.95 |
| n7 | 1031-1253 | 33-55 | AAGCATATTGACGCATACAAAAC | 36 | 1.5 | -31.7 | 75.8 | 76.9 | 1.047 |
| n8 | 1031-1253 | 71-93 | GCCTAAAAAGGACAAAAAGAAGA | 31 | 2 | -31 | 71.7 | 72.1 | 1.039 |
