## Supplementary Table 7 for "A Computational Approach to Design Potential siRNA Molecules as a Prospective Tool for Silencing Nucleocapsid Phosphoprotein and Surface Glycoprotein Gene of SARS-CoV-2"

A Computational Approach to Design Potential siRNA Molecules as a Prospective Tool for Silencing Nucleocapsid Phosphoprotein and Surface Glycoprotein Gene of SARS-CoV-2

Umar Faruq Chowdhury^1^, Mohammad Umer Sharif Shohan^1^, Kazi Injamamul Hoque^1^ Mirza Ashikul Beg^1^, Mohammad Ali Moni^2^, Mohammad Kawsar Sharif Siam^3^

| **Table S7. Effective siRNAs against surface glycoprotein with GC%, free energy of folding and free energy of binding with target.** | | | | | | | | | |
| --- | --- | --- | --- | --- | --- | --- | --- | --- | --- |
| **Alias** | **Conserved  position** | **Location of target  within mRNA** | **siRNA target within mRNA** | **GC %** | **Free energy of folding** | **Free energy  of binding** | **Tm (conc)** | **Tm (Cp)** | **Validity (binary)** |
| g1 | 146-221 | 22-44 | TACCTTTCTTTTCCAATGTTACT | 31 | 1.7 | -29.6 | 78.1 | 79.5 | 0.991 |
| g2 | 223-470 | 88-110 | TGGATTTTTGGTACTACTTTAGA | 29 | 1.8 | -30.1 | 77.8 | 79.2 | 1.031 |
| g3 | 223-470 | 144-166 | CGCTACTAATGTTGTTATTAAAG | 26 | 1.6 | -28.9 | 76.5 | 77.7 | 0.993 |
| g4 | 223-470 | 192-214 | TCCATTTTTGGGTGTTTATTACC | 31 | 2 | -29.5 | 77.3 | 78.1 | 0.986 |
| g5 | 223-470 | 201-223 | GGGTGTTTATTACCACAAAAACA | 31 | 1.8 | -30.9 | 78.2 | 78.7 | 1.038 |
| g6 | 472-541 | 15-37 | TGCGAATAATTGCACTTTTGAAT | 31 | 1.8 | -31.3 | 78.7 | 78.9 | 0.983 |
| g7 | 472-541 | 16-38 | GCGAATAATTGCACTTTTGAATA | 29 | 1.7 | -30.4 | 76.8 | 77.6 | 0.973 |
| g8 | 543-661 | 25-47 | TAGGGAATTTGTGTTTAAGAATA | 29 | 1.8 | -29.7 | 77.7 | 78.9 | 1.026 |
| g9 | 543-661 | 27-49 | GGGAATTTGTGTTTAAGAATATT | 24 | 1.8 | -29.2 | 74.7 | 75.8 | 1.029 |
| g10 | 543-661 | 41-63 | AAGAATATTGATGGTTATTTTAA | 19 | 1.8 | -25.5 | 72.6 | 73.7 | 1.046 |
| g11 | 543-661 | 48-70 | TTGATGGTTATTTTAAAATATAT | 14 | 1.8 | -23.3 | 69.4 | 70.5 | 1.086 |
| g12 | 742-869 | 75-97 | TAGGACTTTTCTATTAAAATATA | 19 | 1.5 | -25.8 | 73.9 | 74.3 | 1.111 |
| g13 | 742-869 | 78-100 | GACTTTTCTATTAAAATATAATG | 14 | 1.5 | -22.9 | 67.4 | 67.7 | 1.004 |
| g14 | 742-869 | 83-105 | TTCTATTAAAATATAATGAAAAT | 10 | 1.6 | -21.6 | 66.4 | 66.3 | 1.012 |
| g15 | 871-1222 | 20-42 | CAGAAACAAAGTGTACGTTGAAA | 36 | 1.6 | -32.8 | 79.1 | 80.2 | 1.022 |
| g16 | 871-1222 | 53-75 | TAGAAAAAGGAATCTATCAAACT | 26 | 1.9 | -27.8 | 75.3 | 76.5 | 1.014 |
| g17 | 871-1222 | 59-81 | AAGGAATCTATCAAACTTCTAAC | 31 | 1.3 | -30.6 | 77.9 | 78.9 | 0.96 |
| g18 | 871-1222 | 60-82 | AGGAATCTATCAAACTTCTAACT | 29 | 1.4 | -30.3 | 76.9 | 78 | 0.955 |
| g19 | 871-1222 | 64-86 | ATCTATCAAACTTCTAACTTTAG | 26 | 1.7 | -27.9 | 75.5 | 76.8 | 0.953 |
| g20 | 871-1222 | 107-129 | TTGTTAGATTTCCTAATATTACA | 21 | 1.8 | -26 | 73.8 | 74.3 | 0.99 |
| g21 | 871-1222 | 136-158 | TGCCCTTTTGGTGAAGTTTTTAA | 36 | 1.8 | -33.4 | 83.4 | 84.6 | 1.039 |
| g22 | 871-1222 | 138-160 | CCCTTTTGGTGAAGTTTTTAACG | 33 | 1.9 | -30.4 | 78.6 | 79.6 | 1.01 |
| g23 | 871-1222 | 162-184 | CACCAGATTTGCATCTGTTTATG | 38 | 2 | -33.7 | 82.3 | 81.1 | 0.902 |
| g24 | 871-1222 | 218-240 | CTGATTATTCTGTCCTATATAAT | 26 | 1.8 | -30.8 | 77 | 78.3 | 1.052 |
| g25 | 871-1222 | 242-264 | CCGCATCATTTTCCACTTTTAAG | 36 | 1.8 | -32.9 | 80.4 | 81.9 | 0.985 |
| g26 | 871-1222 | 243-265 | CGCATCATTTTCCACTTTTAAGT | 31 | 1.8 | -30.5 | 77.9 | 79.2 | 1.021 |

| **Table S7. Effective siRNAs against surface glycoprotein with GC%, free energy of folding and free energy of binding with target. (Continued)** | | | | | | | | | |
| --- | --- | --- | --- | --- | --- | --- | --- | --- | --- |
| **Alias** | **Conserved  position** | **Location of target  within mRNA** | **siRNA target within mRNA** | **GC %** | **Free energy of folding** | **Free energy  of binding** | **Tm (conc)** | **Tm (Cp)** | **Validity (binary)** |
| g27 | 871-1222 | 317-339 | ATGCAGATTCATTTGTAATTAGA | 26 | 1.8 | -28.1 | 75.6 | 76.5 | 1.012 |
| g28 | 1224-1421 | 21-43 | CTGGAAAGATTGCTGATTATAAT | 31 | 1.8 | -32.2 | 79.5 | 80.7 | 1.049 |
| g29 | 1224-1421 | 22-44 | TGGAAAGATTGCTGATTATAATT | 26 | 1.8 | -30.3 | 77.7 | 79 | 1.053 |
| g30 | 1224-1421 | 72-94 | GCGTTATAGCTTGGAATTCTAAC | 36 | 1.8 | -33.8 | 80.7 | 82 | 0.971 |
| g31 | 1224-1421 | 88-110 | TTCTAACAATCTTGATTCTAAGG | 29 | 1.7 | -28.3 | 74.8 | 75.9 | 1.02 |
| g32 | 1224-1421 | 103-125 | TTCTAAGGTTGGTGGTAATTATA | 33 | 1.8 | -32.4 | 82.3 | 83.7 | 0.978 |
| g33 | 1427-1622 | 140-162 | AACTGTTTGTGGACCTAAAAAGT | 36 | 1.8 | -31.6 | 81.4 | 82.8 | 0.931 |
| g34 | 1427-1622 | 151-173 | GACCTAAAAAGTCTACTAATTTG | 29 | 1.9 | -29 | 77.1 | 78.3 | 0.931 |
| g35 | 1427-1622 | 152-174 | ACCTAAAAAGTCTACTAATTTGG | 26 | 1.6 | -27.6 | 75.4 | 76.6 | 1 |
| g36 | 1427-1622 | 159-181 | AAGTCTACTAATTTGGTTAAAAA | 24 | 1.8 | -27.7 | 76.3 | 77.5 | 0.92 |
| g37 | 1427-1622 | 161-183 | GTCTACTAATTTGGTTAAAAACA | 24 | 1.7 | -26.7 | 73.6 | 74.5 | 1.026 |
| g38 | 1624-1840 | 29-51 | TTCTTACTGAGTCTAACAAAAAG | 31 | 1.5 | -30.2 | 78.3 | 79.5 | 0.985 |
| g39 | 1842-2389 | 229-251 | ATCCATCATTGCCTACACTATGT | 40 | 1.7 | -35.3 | 83.5 | 84.8 | 0.92 |
| g40 | 1842-2389 | 406-428 | CAGCAATCTTTTGTTGCAATATG | 33 | 2 | -31 | 79 | 79.8 | 0.931 |
| g41 | 1842-2389 | 419-441 | TTGCAATATGGCAGTTTTTGTAC | 36 | 2 | -31.7 | 80.1 | 80.9 | 0.943 |
| g42 | 1842-2389 | 428-450 | GGCAGTTTTTGTACACAATTAAA | 29 | 1.7 | -31.3 | 77.9 | 79.1 | 1.097 |
| g43 | 1842-2389 | 430-452 | CAGTTTTTGTACACAATTAAACC | 26 | 1.7 | -26.1 | 72.7 | 74 | 1.096 |
| g44 | 1842-2389 | 441-463 | CACAATTAAACCGTGCTTTAACT | 33 | 1.9 | -30 | 79.5 | 80.2 | 1.102 |
| g45 | 1842-2389 | 469-491 | AGCTGTTGAACAAGACAAAAACA | 33 | 1.5 | -31.2 | 79.6 | 80.2 | 0.994 |
| g46 | 1842-2389 | 482-504 | GACAAAAACACCCAAGAAGTTTT | 36 | 1.6 | -32.5 | 80.6 | 81.8 | 1.076 |
| g47 | 1842-2389 | 512-534 | GTCAAACAAATTTACAAAACACC | 26 | 1.7 | -26 | 70.8 | 72 | 0.953 |
| g48 | 1842-2389 | 524-546 | TACAAAACACCACCAATTAAAGA | 31 | 1.8 | -29.5 | 78.5 | 79.9 | 1.045 |
| g49 | 2391-2471 | 3-25 | GTGGTTTTAATTTTTCACAAATA | 21 | 1.6 | -26 | 72.3 | 72.8 | 1.057 |
| g50 | 2391-2471 | 53-75 | AGGTCATTTATTGAAGATCTACT | 29 | 1.7 | -30.5 | 76.4 | 77.3 | 0.97 |
| g51 | 2473-2762 | 226-248 | ATGCAAATGGCTTATAGGTTTAA | 33 | 1.9 | -32.1 | 82.2 | 83.5 | 0.964 |
| g52 | 2473-2762 | 261-283 | TACACAGAATGTTCTCTATGAGA | 36 | 1.9 | -33.2 | 79.5 | 79.7 | 0.935 |
| g53 | 2790-3822 | 68-90 | AACCAAAATGCACAAGCTTTAAA | 33 | 1.9 | -31.5 | 81.6 | 83.3 | 0.961 |
| g54 | 2790-3822 | 69-91 | ACCAAAATGCACAAGCTTTAAAC | 33 | 1.8 | -31.5 | 80.4 | 81.8 | 0.991 |
| g55 | 2790-3822 | 123-145 | GTGCAATTTCAAGTGTTTTAAAT | 26 | 1.8 | -28.7 | 75.6 | 76.9 | 1.055 |
| g56 | 2790-3822 | 124-146 | TGCAATTTCAAGTGTTTTAAATG | 24 | 1.6 | -26.7 | 74.4 | 75.4 | 1.014 |
| g57 | 2790-3822 | 130-152 | TTCAAGTGTTTTAAATGATATCC | 24 | 1.6 | -24.9 | 71.9 | 72.7 | 0.99 |
| g58 | 2790-3822 | 149-171 | ATCCTTTCACGTCTTGACAAAGT | 40 | 1.6 | -33.7 | 82 | 83 | 0.901 |
| g59 | 2790-3822 | 304-326 | GTGTGTACTTGGACAATCAAAAA | 36 | 1.5 | -33.9 | 80.4 | 81.4 | 1.007 |
| g60 | 2790-3822 | 449-471 | GCCATTTGTCATGATGGAAAAGC | 38 | 1.8 | -33.9 | 79.4 | 79.9 | 0.94 |

| **Table S7. Effective siRNAs against surface glycoprotein with GC%, free energy of folding and free energy of binding with target. (Continued)** | | | | | | | | | |
| --- | --- | --- | --- | --- | --- | --- | --- | --- | --- |
| **Alias** | **Conserved  position** | **Location of target  within mRNA** | **siRNA target within mRNA** | **GC %** | **Free energy of folding** | **Free energy  of binding** | **Tm (conc)** | **Tm (Cp)** | **Validity (binary)** |
| g61 | 2790-3822 | 519-541 | TTGTAACACAAAGGAATTTTTAT | 24 | 1.7 | -26.9 | 75.7 | 76.4 | 0.914 |
| g62 | 2790-3822 | 636-658 | AACCTGAATTAGACTCATTCAAG | 36 | 1.7 | -32.7 | 79.5 | 80.4 | 0.984 |
| g63 | 2790-3822 | 649-671 | CTCATTCAAGGAGGAGTTAGATA | 40 | 1.6 | -36.6 | 82.9 | 84.1 | 0.922 |
| g64 | 2790-3822 | 659-681 | GAGGAGTTAGATAAATATTTTAA | 21 | 1.8 | -27.7 | 73.9 | 75.1 | 1.075 |
| g65 | 2790-3822 | 660-682 | AGGAGTTAGATAAATATTTTAAG | 19 | 1.8 | -26.2 | 72.3 | 73.3 | 1.078 |
| g66 | 2790-3822 | 738-760 | TTGTAAACATTCAAAAAGAAATT | 19 | 1.8 | -24.6 | 71.5 | 72.5 | 0.962 |
| g67 | 2790-3822 | 809-831 | CTCCAAGAACTTGGAAAGTATGA | 38 | 1.9 | -33.8 | 82 | 82.4 | 0.928 |
| g68 | 2790-3822 | 820-842 | TGGAAAGTATGAGCAGTATATAA | 31 | 1.8 | -33.1 | 81 | 82.3 | 0.954 |
| g69 | 2790-3822 | 851-873 | TGGTACATTTGGCTAGGTTTTAT | 36 | 1.8 | -34.2 | 83.8 | 85.2 | 0.926 |
| g70 | 2790-3822 | 1001-1023 | GTGCTCAAAGGAGTCAAATTACA | 38 | 1.9 | -33.7 | 81.7 | 82.8 | 1.012 |
