## Supplementary Table 8 for "A Computational Approach to Design Potential siRNA Molecules as a Prospective Tool for Silencing Nucleocapsid Phosphoprotein and Surface Glycoprotein Gene of SARS-CoV-2"

| **Table S8. Bootstrap values of significant clades in the radial phylogenetic tree.** | | | | | | | |
| --- | --- | --- | --- | --- | --- | --- | --- |
| **Significant clades of nucleocapsid phosphoprotein gene** | | | | **Significant clades of surface glycoprotein gene** | | | |
| **Accession** | **Size** | **Country** | **Bootstrap value** | **Accession** | **Size** | **Country** | **Bootstrap value** |
| MT246470 | 29860 | USA | 0.66 | MT039888 | 29882 | USA | 0.62 |
| MT246451 | 29868 | USA |  | MT072688 | 29811 | Nepal |  |
| MT233522 | 29782 | Spain |  | MT044257 | 29882 | USA |  |
| MT246456 | 29846 | USA |  | MN994467 | 29882 | USA |  |
| MT106054 | 29882 | USA | 0.63 | MN988713 | 29882 | USA |  |
| MN997409 | 29882 | USA |  | MT246463 | 29782 | USA | 0.65 |
| MN975262 | 29891 | China |  | MT246456 | 29846 | USA |  |
| MN938384 | 29838 | China |  | MT246482 | 29844 | USA |  |
| MT123291 | 29882 | China | 0.66 | MT246488 | 29842 | USA | 0.86 |
| MT123293 | 29871 | China |  | MT246461 | 29877 | USA |  |
| MT159717 | 29882 | USA | 0.62 | MN996527 | 29825 | China | 0.65 |
| MT184911 | 29882 | USA |  | MT106053 | 29882 | USA |  |
| MT233523 | 29782 | Spain | 0.87 | MT246489 | 29864 | USA | 0.63 |
| MT233519 | 29782 | Spain |  | MT246478 | 29889 | USA |  |

**Supplementary Table 8:**

A Computational Approach to Design Potential siRNA Molecules as a Prospective Tool for Silencing Nucleocapsid Phosphoprotein and Surface Glycoprotein Gene of SARS-CoV-2

Umar Faruq Chowdhury^1^, Mohammad Umer Sharif Shohan^1^, Kazi Injamamul Hoque^1^ Mirza Ashikul Beg^1^, Mohammad Ali Moni^2^, Mohammad Kawsar Sharif Siam^3^
